# mTORC1 and STAT3 signalings are indispensable for in vitro TGFβ1-dependent Three-Dimensional (3D) tendon construct

**DOI:** 10.1101/2024.09.26.615201

**Authors:** Bon-hyeock Koo, Aiden Smith, Kyu Sang Joeng

## Abstract

The Transforming Growth Factor-β 1 (TGFβ1) is a well-known growth factor involved in tenocytes differentiation, extracellular matrix production, and cell fate regulation. We previously demonstrated that TGFβ1 has a critical role in the formation of *in vitro* 3D tendon constructs using mouse primary tendon cells. In this study, we investigated the function of Mammalian target of rapamycin complex 1 (mTORC1) and Signal transducer and activator of transcription 3 (STAT3) signaling in the formation of TGFβ1-induced *in vitro* 3D tendon constructs using specific inhibitors, rapamycin (mTORC1 inhibitor) and stattic (stat3 inhibitor). TGFβ1 treatment activated both mTORC1 and STAT3 in 3D tendon constructs. The treatment of rapamycin or stattic partly attenuated TGFβ1-dependent cellular, molecular, and matrix changes in the 3D tendon constructs. Overall, this study demonstrates that mTORC1-STAT3 signaling axis is a downstream mediator of TGFβ1 signaling in the formation of 3D tendon constructs.

## Introduction

Tendons transmit mechanical forces from muscles to bones, enabling the physical movement of the body [1]. The strength of tendons is derived from a highly organized collagen fiber structure, primarily composed of type I collagen [2]. Tendon fibroblasts, tenocytes, produce and organize the extracellular matrix (ECM) of tendon [3]. Cellular differentiation and morphological maturation of tenocytes are required for proper tendon formation [4]. Scleraxis (Scx) is a critical transcriptional factor regulating tenocyte differentiation by regulating type I collagen expression [5, 6]. Tenomodulin (Tnmd) is a type II transmembrane protein required for tenocyte proliferation and maturation of collagen fibers [7]. Mature tenocytes exhibit elongated, spindle-shaped cell bodies with a reduced number of protrusions [8]. Various growth factors such as TGFβ1, CTGF, BMP1/2, and IGF1 are also involved in tendon development and maturation [9-12].

*In vitro* studies using a monolayer cell culture system contribute to the understanding of molecular mechanisms regulating the development of the musculoskeletal system. However, monolayer tendon cell culture represents several challenges. Primary cells from animal tendon tissues exhibit variability based on the specific tendon region, isolation method, and the age of the animal subject [13, 14]. Tenocytes struggle to maintain their phenotype in monolayer culture, leading to reduced collagen accumulation and adopting a rounder shape rather than their characteristic elongated form as passages progress [15]. These challenges prompted us to develop a reliable scaffold-free 3D *in vitro* tendon culture system, where the peripheral layers of the constructs exhibit tendon-like tissue formation. The constructs underwent tissue maturation process similar to tendon, such as increased tissue thickness, decreased cell density, highly aligned collagen fibers, and elongated cell shapes [16]. Notably, TGFβ1 treatment induced the formation of tendon-like structures, and it increased peripheral layer thickness by maintaining the proliferation and collagen fiber formation in this system. In addition, TGFβ1 induced Scx expression in the tendon constructs [16]. These data suggest that the constructs can be a useful tool for studying the molecular and cellular mechanism regulating tendon development and maturation.

The TGFβ1 is a well-known growth factor mediating various biological processes via SMAD, MAPKs (Mitogen-activated protein kinases), and PI3K (phosphatidyl inositol 3-kinase)-Akt signaling pathway [17, 18]. TGFβ1 signaling is essential for recruiting tendon progenitor cells to tendons and plays a crucial role in neonatal tendon regeneration [19]. Additionally, it contributes to the tendon healing process in the tendon injury model [20]. TGFβ1 superfamily induces the expression of Scx, MMPs, and type I and III collagens in tendon fibroblasts during resistance exercise, triggering matrix turnover and collagen deposition throughout the maturation process [21-23]. Despite the well-known role of TGFβ1 signaling in tendons, the downstream signaling pathways mediating its function remain unclear. Our previous study demonstrated that TGFβ1 induces the expression of the early tenogenic differentiation marker, *Scx*, and enhances the morphological maturation of tendon cells in a 3D tendon model [16]. Therefore, TGFβ1-treated 3D tendon constructs can be utilized to elucidate the downstream mediators of TGFβ1 signaling in tendons.

The mTORC1 (mammalian target of rapamycin complex 1) is a multifaceted signaling pathway involved in cellular proliferation, differentiation, and autophagy regulation, functioning as a downstream mediator of various growth factors [24-26]. Recent mouse genetic studies have demonstrated that both loss and gain of mTORC1 signaling in tendon resulted in impaired postnatal tendon development, suggesting that finely tuned regulation of mTORC1 signaling is critical for this process [27]. Additionally, activation of mTORC1 signaling in tendon stem and progenitor cells is associated with the development of mechanical stress-mediated tendinopathy and age-related tendon disorders [28]. Despite the function of mTORC1 signaling in physiological and pathological tendon conditions, the upstream regulators and downstream mediators of mTORC1 signaling in tendons remain poorly understood.

The STAT3 (signal transducer and activator of transcription 3) is a transcriptional factor that plays a pivotal role in various tissue development and function [29]. The STAT3 signaling pathway is associated with attenuated inflammation and scar formation during tendon healing [30]. In addition, STAT3 phosphorylation is promoted primarily by vascular proliferation in ruptured rotator cuff tendons [31]. Inhibition of the STAT3 signaling pathway reduces ECM generation and fibroblast proliferation from hypertrophic scars [32]. STAT3 can be activated via phosphorylation, with the tyrosine 705 site phosphorylated by JAK [33]. Interestingly, previous cancer studies have demonstrated that mTORC1 can activate STAT3 via phosphorylation of serine-727 (S727) [34]. However, the interaction between STAT3 and mTORC1 signaling in tendons has not been established.

In this study, we investigated the role of the mTORC1-STAT3 signaling axis in TGFβ1-induced scaffold-free 3D tendon constructs. Rapamycin and stattic were employed as inhibitors of mTORC1 and STAT3, respectively, to explore their involvement in TGFβ1-induced tendon-like tissue formation. The findings in current study highlight the pivotal role of the mTORC1-STAT3 axis in regulating tendon thickness, cell proliferation, collagen fibrillogenesis, and molecular changes, such as Scx expression. TGFβ1 treatment in 3D in vitro tendons activates the mTORC1-STAT3 signaling pathway. Overall, these data demonstrate that the mTORC1-STAT3 axis mediates the function of TGFβ1 in 3D tendon formation.

## Materials and Methods

### Animals

University Laboratory Animal Resources (ULAR) at the University of Pennsylvania (Philadelphia, Pennsylvania, USA) and Institutional Animal Care and Use Committee (IACUC) approved all procedures and experiments in this research. The study was complied with the ARRIVE guidelines, and mice were euthanized by carbon dioxide gas asphyxiation using IACUC-reviewed and -approved procedures. The 3D tendon constructs were generated by using tail tendon cells from both male and female mice.

### Reagents

Rapamycin (Cat.No. R8781-200UL), type I collagenase (Cat. No. C0130), and Fetal bovine serum (FBS, Cat. No. F8317) were purchased from Sigma-Aldrich (St. Louis, MO). TGFβ1 recombinant protein (Cat. No. 240-B) was purchased from R&D system (Minneapolis, MN), and stattic (Cat. No. ab120952) was purchased from Abcam (Cambridge, MA). α-MEM (Cat. No. 12571063), Penicillin-Streptomycin (P/S, Cat. No. 15140122), L-Glutamine (Cat. No. 25030081), Human plasma-derived fibronectin (Cat. No. 33016015), and antisera against p-mTORC1 (T2446, Cat. No. PA5-104899) were purchased from Thermo Fisher Scientific (Waltham, MA). Antisera against p-mTORC1 (S2448, Cat. No. #5536), Total-mTORC1 (Cat. No. #2972), p-STAT3 (S727, Cat. No. D1B2J, and Y705, Cat. No. D3A7), Total STAT3 (Cat. No. #9132), p-S6 (Cat. No. #2211), Total-S6 (Cat. No. #2217), α-tubulin (Cat. No. #2144) were purchased from Cell Signaling Technology (Beverly, MA).

### Generation of three-dimensional (3D) tendon constructs

Detailed 3D tendon culture was previously described [16]. Briefly, the tendons from mice tail were isolated at 28 days of age and digested in type I collagenase solution (2mg/mL in 1X PBS, Sigma C0130) by inverting every 10 minutes for one hour in a 37 °C incubator. The tenocytes from digested tendons were cultured in growth media (20% FBS, 1% P/S, 2 mM L-Glutamine in αMEM). To make the growth channel for 3D tendon constructs, we placed the 3D-printed mold into agarose solution on a 6-well plate. The mold was removed from the agarose 30 minutes later, and 3D-printed anchors were inserted into each end of the growth channel. The agarose channel was sterilized for 30 minutes by UV, and then growth area was coated with Human plasma-derived fibronectin (0.375mg/ml in 1X PBS). Tendon cells from culture plates were seeded (2.5 × 10^6^ cells) into each fibronectin-coated growth channel and were condensed for 10 minutes in a 37 °C incubator. Growth media (10% FBS, 1X P/S, 2 mM L-Glutamine, 50 µg/ml L-Ascorbic acid in α-MEM) was added after condensation and these constructs were cultured in a standard cell culture incubator (37 °C, 5% CO_2_, and 95% relative humidity). TGFβ1 (5 ng/ml in 4mM HCL containing 1mg/ml bovine serum albumin) or vehicle (4mM HCL containing 1mg/ml bovine serum albumin) was treated in growth media two days after seeing. The growth media treated with TGFβ1 is called differentiation media. Rapamycin (100 nM) and stattic (10 µM) were treated into the differentiation media 7 days later after TGFβ1 treatment. Differentiation media was replaced every other day till 14 days, and rapamycin and stattic were added in the differentiation media as needed. BrdU (5 µg/ml) was added in the media 2 hours before harvesting the constructs for proliferation assay. All constructs were harvested 14 days after culturing in the differentiation media.

### Western blotting analysis

The tendon constructs were washed with PBS twice and homogenized using a pestle (Cat. No. KT749520-0000, VWR) in RIPA buffer. After quantification using the Quick Start™ Bradford Protein Assay kit (Cat. No. #5000201, Bio-RaD), the lysate was centrifuged at 1,000 *g* for 5 minutes and supernatant was retrieved. The protein lysate was added with 4X Laemmli Sample Buffer (Cat. No. #1610747, Bio-RaD) and boiled at 100 °C for 10 minutes. The prepared samples were used for SDS-polyacrylamide gel electrophoresis, and then the proteins were transferred to polyvinylidene difluoride (PVDF, Cat. No. IPVH00010, Millipore) membrane. The membrane was blocked with EveryBlot Blocking Buffer (Cat. No. 12010020, Bio-RaD) for 1 hour at room temperature, and then incubated with primary antibody (1: 1000) at 4 °C for overnight. After primary antibody incubation, horseradish peroxidase-conjugated secondary antibody was incubated at room temperature for 2 hours. Each protein was detected using ChemiDoc XRS+ System (Hercules, California, Bio-RaD).

### RT-PCR and quantitative real-time PCR (qRT-PCR)

Total RNA from tendon constructs was isolated with Trizol (Cat. No. 15596026, Thermo Fisher Scientific) and Direct-zol RNA miniprep kits (Zymo Research, R2050), and then quantified using NanoDrop ND-1000 Spectrophotometer (Waltham, MA, Thermo Fisher Scientific). cDNA was produced using iScript Reverse Transcription (Bio-Rad, 1708841) from 500 ng of total RNA. The SYBR green master mix (Pec, #4385612) was used for qRT-PCR which was performed on Quantstudio 6 (Waltham, MA, Thermo Fisher Scientific). There are specific primer sequences in Table 1.

**Table 1.**
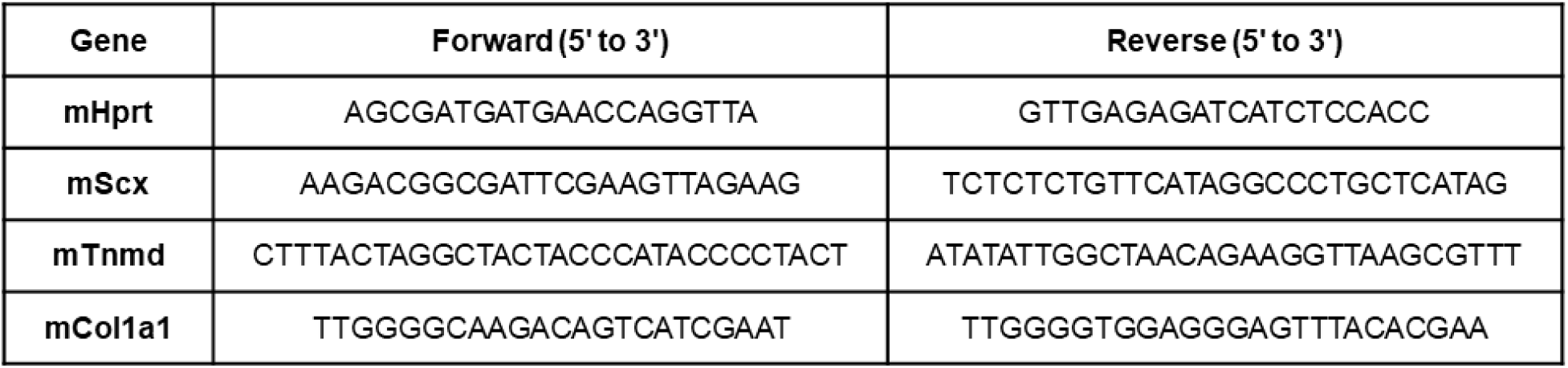
Primer sequences for qRT-PCR.

### Histological analyses

The tendon constructs were imaged on optical microscope to check the status of constructs, before harvesting. The 3D tendons were washed with PBS twice and fixed with 4% paraformaldehyde solution (Cat. No. AAJ19943K2, Thermo Fisher Scientific) at 4 °C for overnight. These samples were embedded in paraffin and sectioned with 6 µm thickness using a microtome. In Hematoxylin and Eosin (H&E) staining, Hematoxylin was stained for 3 minutes after hydrolysis of section samples and Eosin was counterstained for 2 minutes. BrdU staining was performed using BrdU Immunohistochemistry Kit (Cat. No. ab125306, Abcam) and TUNEL assay was performed using TUNEL Assay Kit - HRP-DAB (Cat. No. ab206386). Nucleus aspect ratio was performed by measuring the length between height and width of every single nucleus from the H&E-stained images. The width is always parallel to channel length, and the bigger value indicates more elongated nucleus shape.

### Second Harmonic Generation

The paraffin sections were washed with Xylene twice for 3 minutes and gradually hydrolyzed with ethanol and PBS. After mounting with anti-fade DAPI solution (Invitrogen, P36935), Second Harmonic Generation (SHG) Images were acquired using a Coherent Chameleon Vision II Ti-Sapphire laser (Coherent, Santa Clara, CA) in a Leica TCS SP8 MP multiphoton microscope (Leica Microsystems, Buffalo Grove, IL). The objective lens was a 40x/1.3 NA HC PL APO CS2 oil immersion lens. The images were taken with the 600 Hz scan speed, bidirectional scanning at 880 nm excitation.

### Electron Microscopy Analysis for Collagen Fibril

Harvested 3D tendon constructs were washed with PBS twice and fixed in fixative solution (1.5% glutaraldehyde/1.5% paraformaldehyde with 0.05% tannic acid in DPBS) for overnight in 4 °C cold room. The fixed constructs were performed post-fixation in 1% OsO4 and washed in DPBS. After the samples were dehydrated in ethanol gradually, samples were rinsed in propylene oxide, and Spurrs epoxy infiltrated, and polymerized for freezing at -70 °C overnight. Transmission electron microscopy (TEM) images were acquired from A FEI G20 TEM which shows collagen fibril sections. The length of every single collagen fibril diameter was measured through ImageJ software ([Fiji 2.15.0](https://imagej.net/software/fiji/)). All images were analyzed using five constructs per group, and 200 values were acquired from every single image. ZEISS ZEN software (Carl Zeiss Microscopy, [ZEN 3.9](https://www.zeiss.com/microscopy/en/products/software/zeiss-zen.html#accordionItem-1748894843), Jena, Germany) were used to measure the intensity of the SHG signal. The angle of collagen fibers to the horizontal line were calculated to determine collagen alignment, utilizing ImageJ software. All quantification were performed in a single-blinded manner to ensure unbiased selection.

### Statistical analysis

Results are shown as mean ± SD through dot-graphs. All of experiments were analyzed by one-way ANOVA and conducted using at least three 3D tendon constructs per group. P < 0.01 is considered significant.

## Results

### The formation of TGFβ1-induced 3D tendon constructs was impsired by inhibition of mTORC1-STAT3 signaling pathway

We previously reported that TGFβ1 treatment induced tendon-like tissue structure in the peripheral layer of 3D tendon constructs [16]. To test if mTORC1-STAT3 signaling is involved in TGFβ1-dependent tendon-like tissue formation in tendon constructs, we treated rapamycin (mTORC1 inhibitor) and stattic (STAT3 inhibitor) in the 3D tendon constructs. The treatment timeline is summarized in Figure 1. TGFβ1-treated constructs are thicker and contain fewer budding cells on the surface when compared with vehicle-treated constructs (Figure 2A, first and second panels). Co-treatment of rapamycin or stattic with TGFβ1 reduced the thickness of the tendon constructs compared to TGFβ1-treated constructs (Figure 2A, third and fourth panels). The histological analysis using H&E-stained sections confirmed the reduced thickness with rapamycin or stattic treatment (Figure 2B). More specifically, the TGFβ1-treated constructs maintained greater thickness in both peripheral and inner layers compared to vehicle-treated constructs (Figure 2B, first and second panels, peripheral: p< 0.001, inner: p=0.002). The constructs co-treated with rapamycin or stattic showed reduced thickness in both layers when compared to TGFβ1-treated constructs (Figure 2B second, third, and fourth panels, TGFβ1 vs. TGFβ1 + rapamycin: peripheral p= 0.014, inner p=0.032, TGFβ1 vs. TGFβ1 + stattic: peripheral p= 0.007, inner p=0.007). The quantification analysis of the histological image verified the thickness changes in each treatment group (Figure 2C, D). Low cell density is one of the major features of matured tendons [35-37], and TGFβ1-treated constructs showed decreased cell density in the peripheral layer when compared to the vehicle-treated group (Figure 2E, p< 0.001). Rapamycin or stattic treatment significantly impeded TGFβ1-induced low cell density phenotype in the peripheral layer (Figure 2E, TGFβ1 vs. TGFβ1 + rapamycin, p=0.003, TGFβ1 vs. TGFβ1 + stattic, p=0.011). On the other hand, TGFβ1-induced high cell density phenotype in the inner layer of constructs was not affected by rapamycin or stattic treatment (Figure 2F). Given the result of cell density in the peripheral layer of 3D tendon constructs, we investigated cellular proliferation and apoptosis in both layers by performing BrdU staining and TUNEL assay (Figure 3A, B). Consistent with the previous study, the quantification analysis showed that TGFβ1 treatment increased cellular proliferation, which is evident with more BrdU-positive cells (Figure 3C, peripheral p<0.001, inner p<0.019). Rapamycin or stattic treatment significantly reduced TGFβ1-induced cellular proliferation in peripheral layer (Figure 3C, TGFβ1 vs. TGFβ1 + rapamycin, p=0.011, TGFβ1 vs. TGFβ1 + stattic, p=0.007). TGFβ1 treatment increased cellular proliferation in inner layer, but rapamycin or stattic treatment had no effect on this inner layer phenotype (Figure 3D). Apoptosis levels were not changed in the peripheral layer of each treated group (Figure 3E). TGFβ1 treatment increased apoptosis level in inner layer, but rapamycin and stattic treatment has no effect on this increased apoptosis (Figure 3F, p=0.004). Overall, these results suggest that mTORC1-STAT3 signaling partly mediates TGFβ1-dependent tendon-like tissue formation in the peripheral layer of 3D tendon constructs.

**Figure 1.**
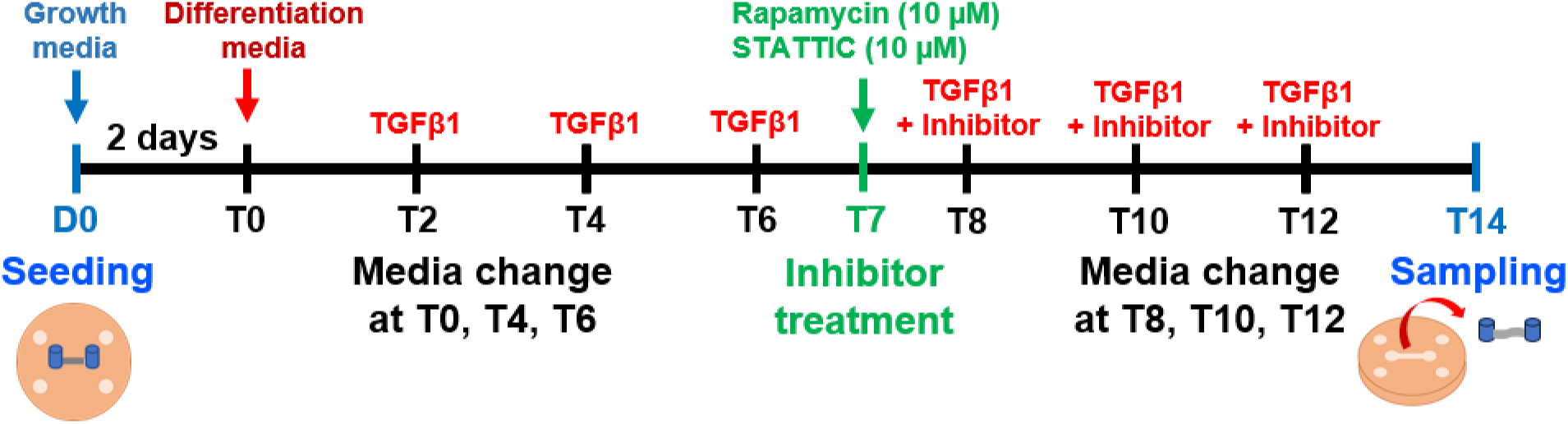
Timeline of 3D tendon culture. After seeding tenocytes from 2D plates, those 3D tendon constructs were cultured in growth media (10% FBS, 1X P/S, 2 mM L-glutamine, and 50 µg/ml L-ascorbic acid) for 2 days. To create constructs for TGFβ1-treated and rapamycin or stattic co-treated groups, the media was changed to differentiation media (10% FBS, 1X P/S, 2 mM L-glutamine, and 50 µg/ml L-ascorbic acid, 5 ng/ml TGFβ1) which time point is T0. The differentiation media was changed every other day (T2, T4, T6). Rapamycin or stattic was co-treated with TGFβ1 at T7. Each inhibitor was co-treated with TGFβ1 every other day (T8, 10, 12) until sampling.

**Figure 2.**
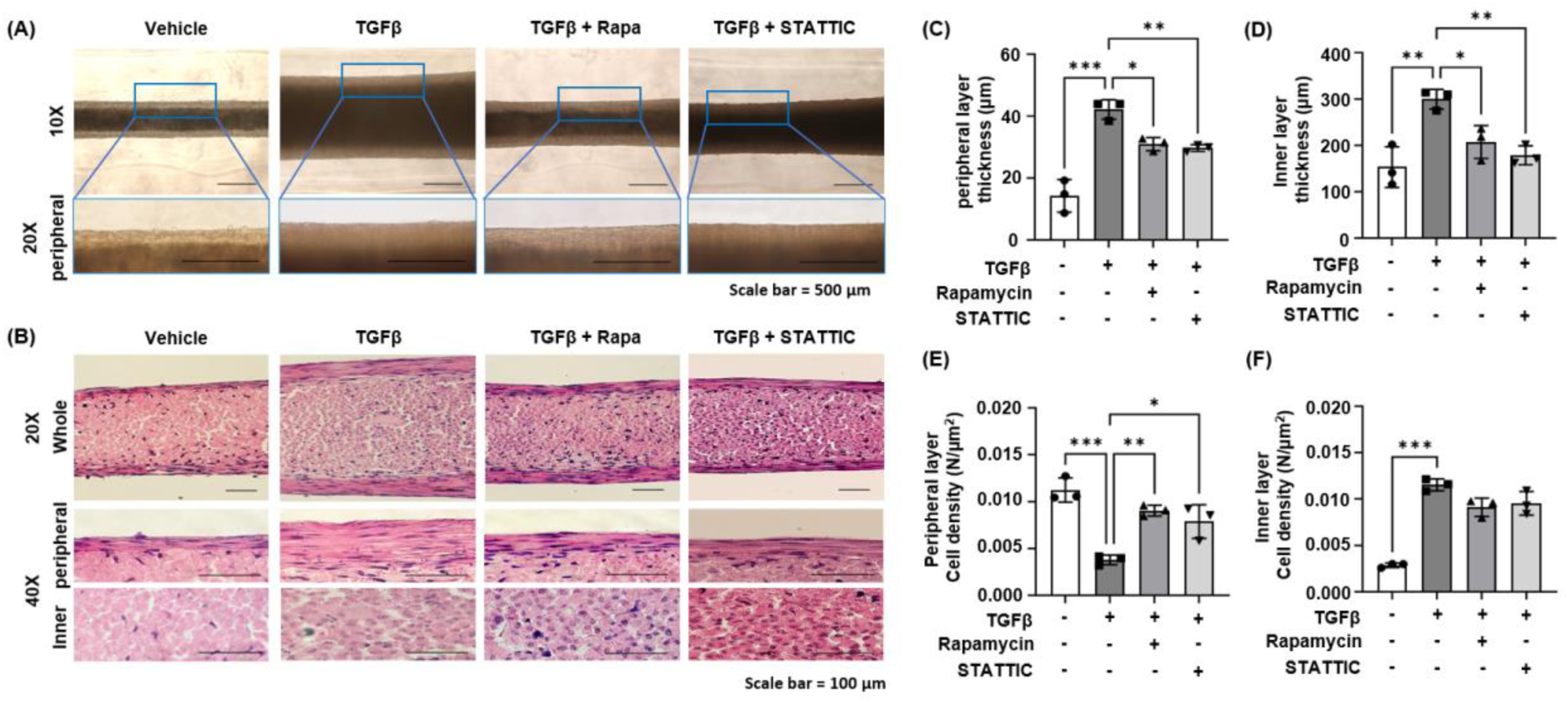
Inhibition of mTORC1-STAT3 signaling impedes tendon-like structural formation. (A) Representative images of 3D live tendon constructs from optical microscope. All groups’ images were acquired at T14. The scale bars indicate 500 µm for both 10X and 20X peripheral images. (B) Representative images of H&E-stained longitudinal sections in whole, peripheral, and inner layer of 3D tendons from vehicle, TGFβ1, TGFβ1 co-treated with rapamycin and stattic at T14. The scale bars indicate 100 µm for both 20X and 40X images. (C, D) Quantification graph of peripheral and inner layer thickness, respectively. (E, F) Quantification graph of peripheral and inner layer cell density, respectively. *** vs. non-treated, p<0.0001, and # vs. TGFβ1, p<0.01, ## vs. TGFβ1, p<0.001 by One way ANOVA from n=3 experiments.

**Figure 3.**
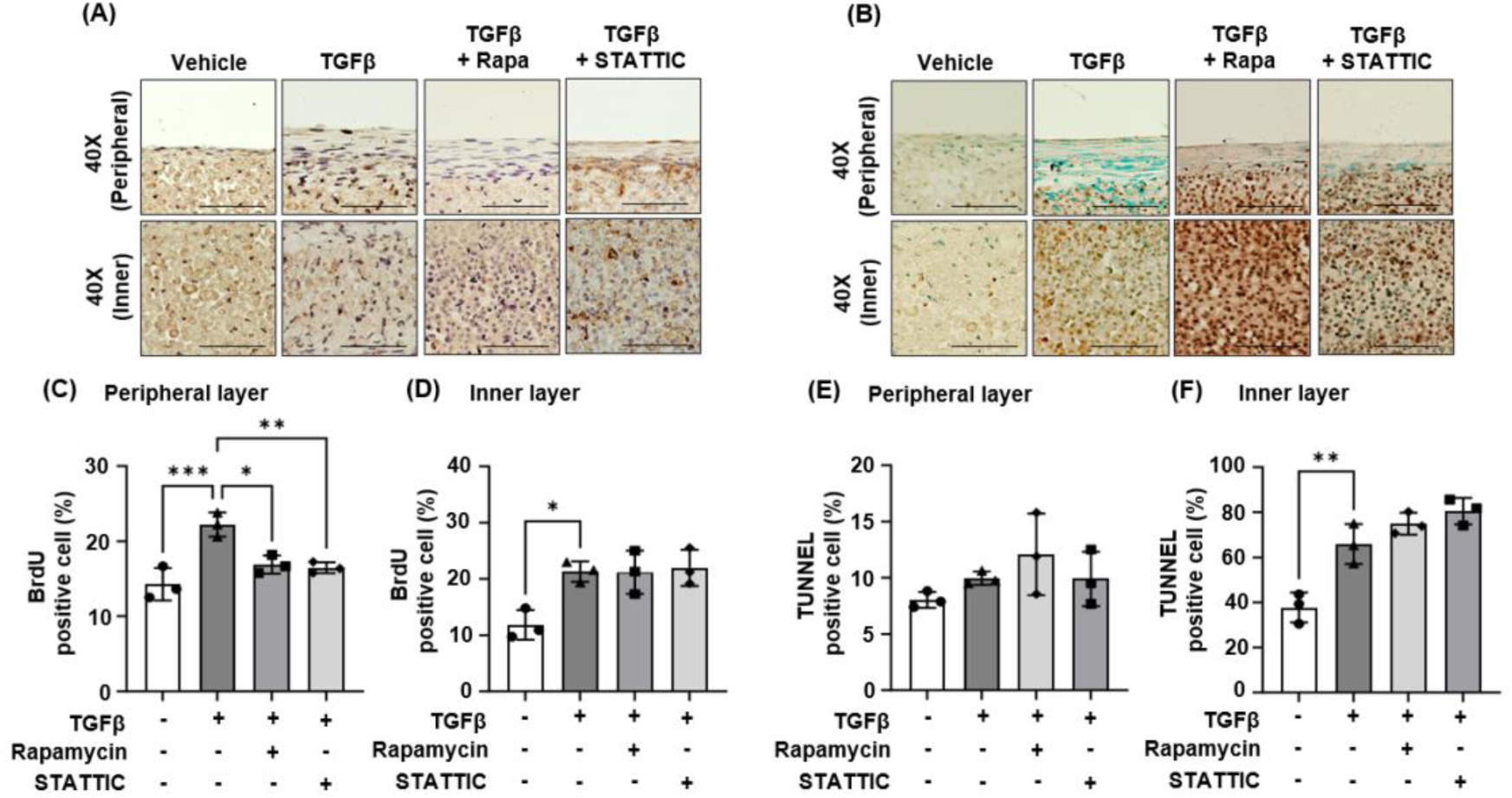
TGFβ1-dependent mTORC1-STAT3 signaling axis up-regulates cellular proliferation in peripheral layer but not apoptosis. (A) Representative images of BrdU stained longitudinal sections in peripheral, and inner layer of 3D tendons. The blue color indicates the single nucleus and brown color indicates the BrdU positive nucleus. (B) Representative images of longitudinal sections from TUNEL assay in peripheral, and inner layer of 3D tendons. The cyan color indicates the single nucleus and brown color indicates the TUNEL positive cells. All groups’ images were acquired at T14. The scale bars indicate 100 µm for both 20X and 40X images. (C, D) Quantification graph of peripheral and inner ratio for BrdU positive cells, respectively. (E, F) Quantification graph of peripheral and inner ratio for TUNEL positive cells, respectively. * vs. non-treated, p<0.01, *** vs. non-treated, p<0.0001, and # vs. TGFβ1, p<0.01 by One way ANOVA from n=3 experiments.

### The treatment of rapamycin or stattic partly inhibited TGFβ1-induced collagen fibrillogenesis and fiber formation in the tendon constructs

Previously, we found that TGFβ1 treatment inhibits the growth of collagen fibril diameter in the tendon constructs [16]. Consistently, transmission electron microscopy (TEM) image showed that TGFβ1-treated constructs exhibited smaller collagen fibril diameter than those of vehicle-treated constructs (Figure 4A). The constructs with rapamycin or stattic treatment revealed increased collagen fibrillar diameter when compared to TGFβ1-treated constructs, but smaller diameter when compared to vehicle-treated constructs (Figure 4A). The quantification results indicated that the major proportion of collagen fibril diameter was 40 nm in vehicle-treated constructs but 30 nm in TGFβ1-treated constructs (Figure 4B, blue and white bar, asterisk). The rapamycin- or stattic-treated group showed the major proportion of collagen fibril at 35 nm (Figure 4B, green and red bar, asterisk). To evaluate the collagen fiber formation and alignment in the peripheral layer, we performed Second Harmonic Generation (SHG) microscopic imaging (Figure 4C). As expected, TGFβ1-treated constructs contain stronger SHG signals than those of vehicle-treated constructs, indicating a higher level of collagen fiber formation (Figure 4C, p<0.001). Treatment of rapamycin or stattic significantly reduced the TGFβ1-induced SHG signal (Figure 4C, TGFβ1 vs. TGFβ1 + rapamycin, p<0.001, TGFβ1 vs. TGFβ1 + stattic, p<0.001), which is verified by the quantification of the fluorescence signal intensity (Figure 4D). The quantification of collagen alignment also demonstrated that TGFβ1 treatment induced more aligned collagen matrix than vehicle-treated constructs, and rapamycin or stattic treatment repressed the TGFβ1-dependent collagen alignment (Figure 4E, - vs. TGFβ1, p<0.001, TGFβ1 vs. TGFβ1 + rapamycin, p<0.001, TGFβ1 vs. TGFβ1 + stattic, p<0.001). These results suggest that the mTORC1-STAT3 signaling axis is partly involved in TGFβ1-dependent collagen fibrillogenesis and fiber formation.

**Figure 4.**
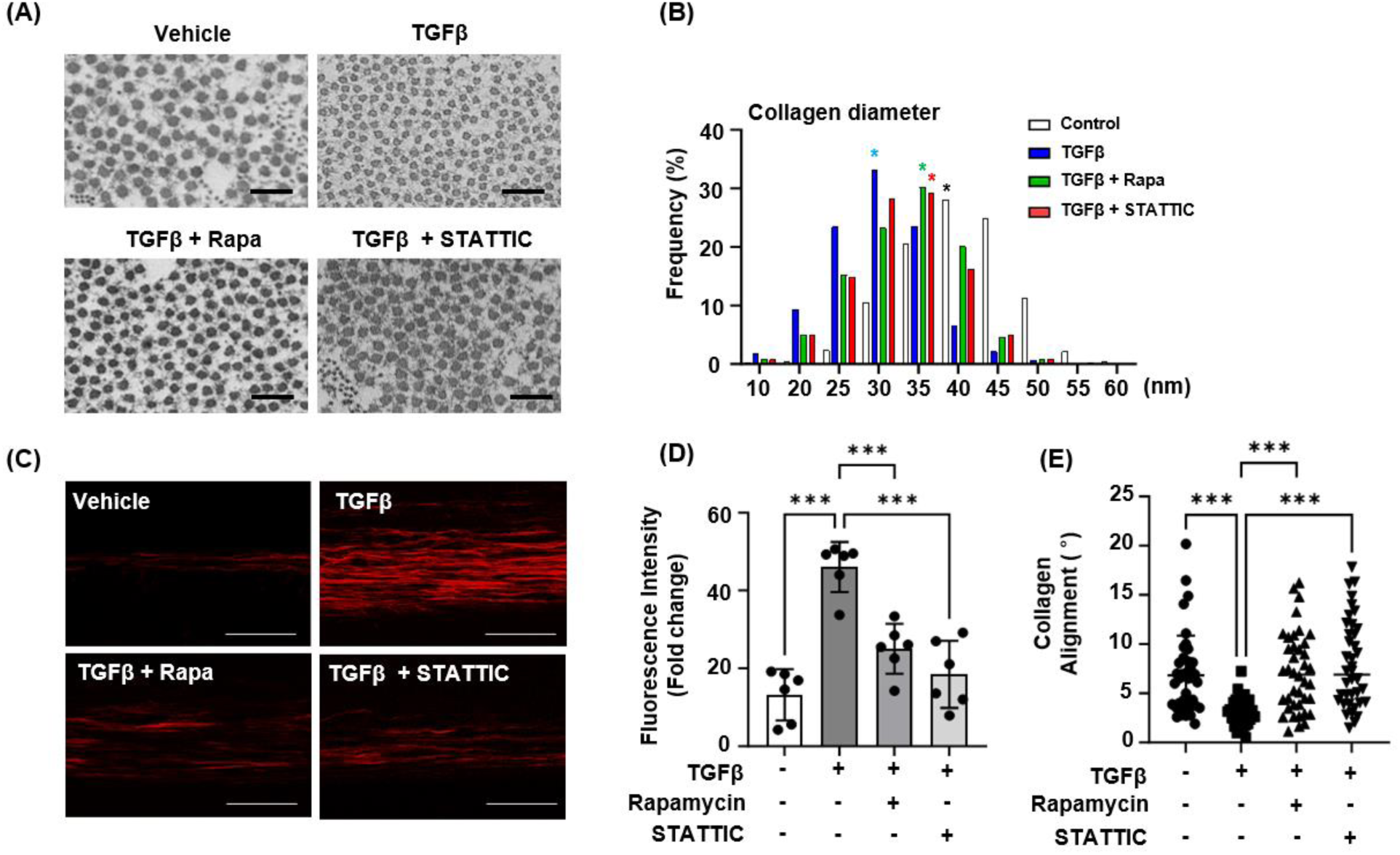
mTORC1-STAT3 signaling determines collagen fibrillogenesis, fiber formation and alignment. (A). Transmission electron microscopic (TEM) images of vehicle, TGFβ1, TGFβ1 co-treated with rapamycin and stattic in peripheral layer at T14. The scale bars indicate 500 nm. (B) Quantification result of each group’s collagen diameter using TEM images at T14 (n=3 experiments). Asterisk mark (*) indicates the highest frequency values in each group. * Vehicle = 40 nm, TGFβ1 = 30 nm, TGFβ1 + rapamycin = 35 nm, TGFβ1 + stattic = 35 nm. (C) Second harmonic generation (SHG) microscopic images for examination of collagen fiber formation and alignment of above groups in peripheral layer at T14. The scale bars indicate 100 µm. (D) The quantification result of collagen fiber formation by measuring fluorescence intensity (red color). *** vs. non-treated, p<0.0001, and ## vs. TGFβ1, p<0.001 by One way ANOVA from n=6 experiments. (E) The quantification result of collagen fiber alignment by measuring the angles of fibers from the basement line. The zero value means the fiber is perfectly parallel to the baseline. *** vs. non-treated, p<0.0001, and ### vs. TGFβ1, p<0.0001 by One way ANOVA from n=6 experiments (The measurement was performed from 40 fibers).

### mTORC1-STAT3 signaling partially mediates the function of TGFβ1 in molecular and morphological changes of cells in the 3D tendon constructs

To investigate the function of mTORC1-STAT3 signaling in TGFβ1-dependent tenogenic differentiation, we performed qRT-PCR with three tenogenic gene markers, *Scleraxis (Scx)*, *Tenomodulin (Tnmd)*, and Collagen type 1 (*Col1a1*) using 3D tendon constructs. Consistent with our previous study, TGFβ1 treatment significantly increased the Scx expression in the tendon constructs (Figure 5A, p<0.001). Rapamycin or stattic treatment significantly reduced the TGFβ1-induced Scx expression (Figure 5A, TGFβ1 vs. TGFβ1 + rapamycin, p<0.001, TGFβ1 vs. TGFβ1 + stattic, p<0.001). Although Tnmd was significantly reduced by TGFβ1 treatment, rapamycin or stattic treatment did not affect the TGFβ1-dependent changes in Tnmd expression (Figure 5B, - vs. TGFβ1, p<0.001). Col1a1 expression was not significantly different between all treated groups (Figure 5C). To examine the morphological maturation of tenocytes in the tendon constructs, we analyzed the nucleus aspect ratio by measuring transverse and perpendicular length in both layers. A value closer to 1.0 would become a circle and more elongated nucleus has greater value. Thus, the distribution of value indicates how many cells would be morphologically changed in the tendon constructs. In the peripheral layer, TGFβ1-treated constructs (average: 5.88) contain more elongated cells than vehicle-treated constructs (average: 3.85), which is indicated by increased nuclear aspect ratio (Figure 5D, p<0.001). Rapamycin- (average: 4.80) or stattic-(average: 4.60) treated group showed reduced nuclear aspect ratio than the TGFβ1-treated group, which indicates less elongated cells in the peripheral layers of these constructs than TGFβ1-treated constructs (Figure 5D, TGFβ1 vs. TGFβ1 + rapamycin, p<0.001, TGFβ1 vs. TGFβ1 + stattic, p<0.001). On the other hand, all treatment groups showed similar frequency of nucleus aspect ratio in inner layer (Figure 5E). These results suggest that mTORC1-STAT3 signaling partially mediates TGFβ1-dependent Scx expression and morphological maturation of tenocytes in the peripheral layer of the tendon constructs.

**Figure 5.**
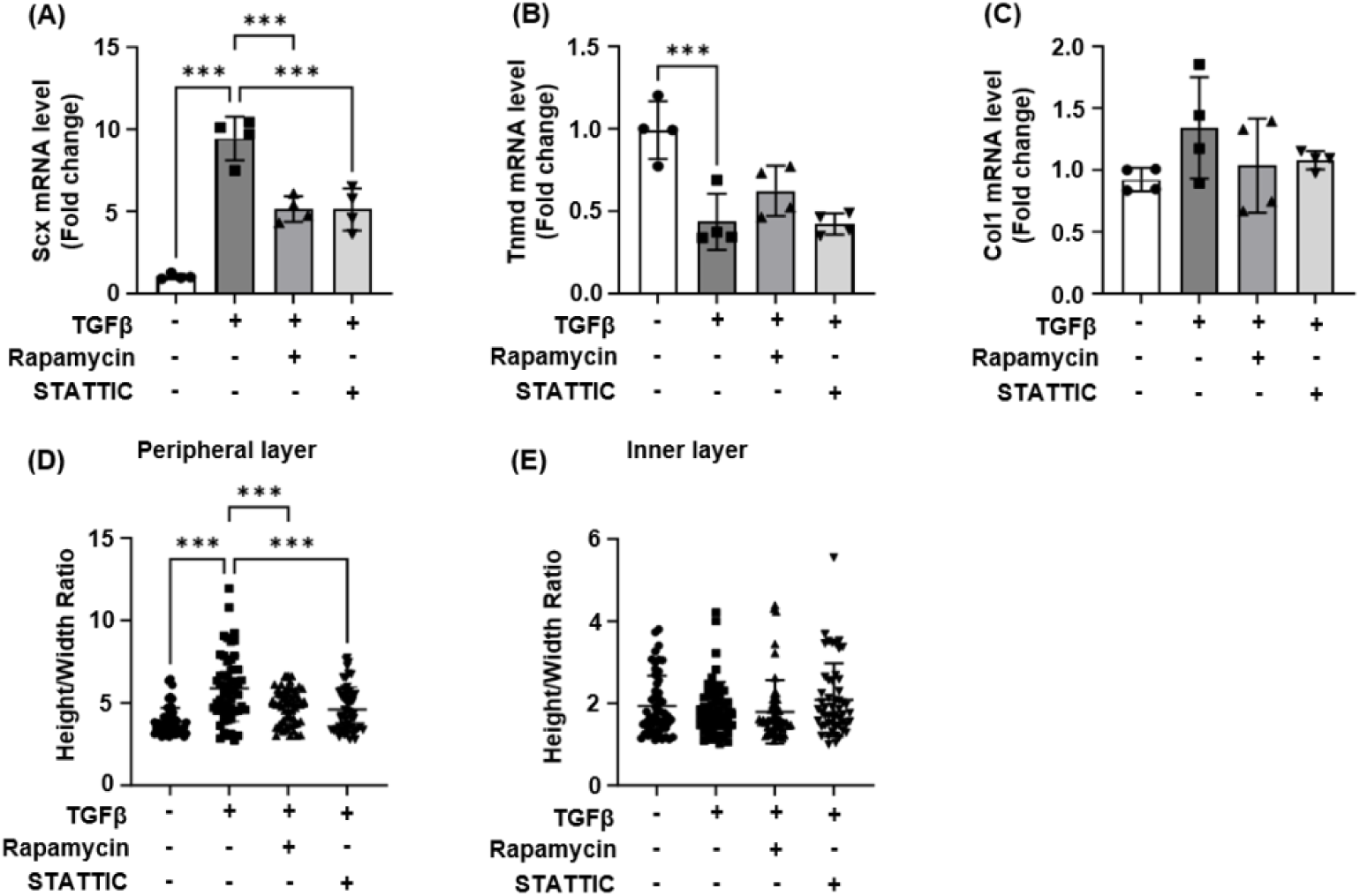
Morphological and molecular changes are induced TGFβ1-dependent mTORC1-STAT3 signaling pathway. (A-C) Quantitative real-time PCR (qRT-PCR) results for tenogenic differentiation markers as Scleraxis (Scx), Tenomodulin (Tnmd), and Type I Collagen (Col1), respectively, in 3D tendon constructs at T14. *** vs. non-treated, p<0.0001, and ### vs. TGFβ1, p<0.0001 by One way ANOVA from n=4 experiments. (D, E) The quantification result of frequency of nucleus aspect ratio by measuring the transverse length to its channel direction length of single nucleus in peripheral layer and inner layer, respectively. The values closer to 1.0 indicates a rounder nucleus shape. n=3 experiments at T14.

### TGFβ1 activates mTOR and STAT3 in 3D tendon constructs

To test whether TGFβ1 activates mTOR and STAT3 in 3D tendon constructs, we examined the phosphorylation of mTOR and STAT3 following TGFβ1 treatment. TGFβ1 treatment increased the phosphorylation of mTOR at the S2448 and T2446 residues (Figure 6A), as confirmed by quantification (Figure 6B, S2448: p<0.001, T2446: p=0.005). Additionally, TGFβ1 treatment enhanced the phosphorylation of STAT3 at the S727 and Y705 residues (Figure 6A), with quantification verifying this increase (Figure 6C, S727: p<0.001, Y705: p=0.004). To further elucidate the TGFβ1-dependent mTOR and STAT3 signaling axis, we investigated the effects of rapamycin and stattic on mTOR and STAT3 phosphorylation in TGFβ1-treated constructs. Rapamycin treatment effectively reduced TGFβ1-induced mTOR phosphorylation, whereas stattic had no effect on mTOR activation (Figure 6B, S2448: p<0.001, T2446: p=0.005). TGFβ1-induced STAT3 phosphorylation was significantly decreased by both rapamycin and stattic (Figure 6C, S727: TGFβ1 vs. TGFβ1 + rapamycin, p<0.001, TGFβ1 vs. TGFβ1 + stattic, p<0.001, Y705: TGFβ1 vs. TGFβ1 + rapamycin, p<0.001, TGFβ1 vs. TGFβ1 + stattic, p<0.001). These findings suggest that mTOR and STAT3 are downstream targets of TGFβ1 in the tendon constructs and that STAT3 is further downstream of mTOR signaling.

**Figure 6.**
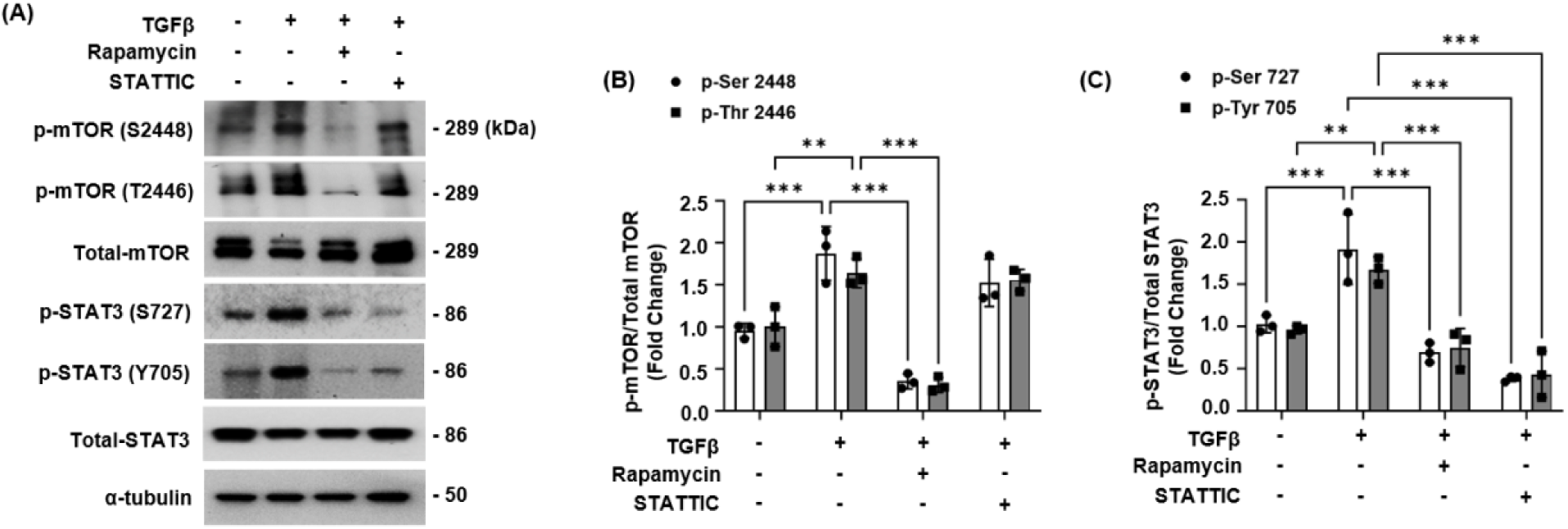
Activation of mTORC-STAT3 signaling axis by TGFβ1 treatment. (A) Western blotting analysis was conducted using 3D tendon constructs of above groups. TGFβ1 (5 ng/ml, 14 days), rapamycin (100 nM, 7 days), and stattic (10 µM, 7 days) were incubated in each group. These constructs were harvested at T14 time point (T means the days after treating with TGFβ1). The α-tubulin served as the control indicating total protein amount of each sample. (B) Quantification group for p-mTOR (p-Ser2448, p-Thr 2446) per total-mTOR. *** vs. non-treated group, p< 0.0001 by One-Way ANOVA test from n=3 experiments. (C) Quantification graph for p-STAT3 (p-Ser727, Tyr-705) per total-STAT3. *** vs. non-treated, p<0.0001, and ### vs. TGFβ1, p<0.0001 by One way ANOVA from n=3 experiments.

## Discussion

In this study, we explored the role of mTORC1 and STAT3 signaling in the formation of TGFβ1-dependent scaffold-free, three-dimensional tendon constructs. Inhibition of the mTORC1 or STAT3 signaling using rapamycin or stattic hindered the TGFβ1-induced tendon-like structure in the peripheral layer of constructs. Treatment of rapamycin or stattic partially impeded TGFβ1-induced collagen fibrillogenesis and collagen fiber formation in the constructs. Moreover, both treatments inhibited TGFβ1-dependent *Scx* expression and morphological maturation of tenocytes. Our findings also revealed that TGFβ1 activates mTORC1 and STAT3 in the constructs. Overall, this is the first *in vitro* study to demonstrate that mTORC1 and STAT3 are downstream mediators of TGFβ1 signaling during the formation of 3D tendon constructs.

Our findings support an interaction between TGFβ1 and mTORC1 in regulating 3D tendon formation. Previous studies have shown similar interactions in pathological conditions. For example, TGFβ1 activates mTORC1 signaling via PDGFRβ to induce mesangial cell hypertrophy and matrix protein accumulation [38]. Additionally, TGFβ1 signaling enhances the expression of PI3K, Akt, and mTOR in pathological scar fibroblasts [39]. mTORC1 also regulates TGFβ1-induced epithelial-mesenchymal transition by reducing pyruvate kinase expression in cervical cancer cells [40]. These findings align with the current study, demonstrating that mTORC1 is a downstream mediator of TGFβ1 signaling in 3D tendon formation.

The current study also suggests that STAT3 is essential for TGFβ1-induced 3D tendon formation. Previous research provides strong evidence that STAT3 mediates various biological processes regulated by TGFβ1 signaling. STAT3 activity and expression are modulated by multiple kinases, some of which are activated by non-canonical TGFβ1 signaling pathways [41]. Inhibition of STAT3 activity has been shown to attenuate TGFβ1-induced fibroblast-to-myofibroblast transition and collagen release [42]. Additionally, TGFβ directly activates STAT3 via JAK1 and SMAD regulation in hepatic cells to control cytokine and growth factor expression [43]. Studies using pharmacological inhibitors and genetic mouse models have also shown STAT3’s partial involvement in TGFβ-mediated fibrosis [42]. We provide further evidence that STAT3 acts as a downstream mediator of TGFβ1 signaling during the molecular and cellular changes involved in 3D tendon formation.

Although our study indicates that mTORC1 and STAT3 function downstream of TGFβ1 during 3D tendon formation, the detailed interaction between STAT3 and mTORC1 remains unclear. Other studies have reported interactions between these pathways in various biological systems. mTORC1 is linked to JAK-STAT3 signaling, and inhibition of mTORC1 activity has been shown to reduce phosphorylation of STAT3 at S727 and Y705 in IL-6-dependent mTOR/STAT3 hyperactivation [44]. mTORC1 also drives HIF-1α transcription via STAT3 during tumor vascularization [45]. Yokogami et al. demonstrated a relationship between mTORC1 and STAT3, investigating the phosphorylation of a C-terminal STAT3 peptide by mTOR kinase [46]. In this study, TGFβ1-induced STAT3 phosphorylation was significantly decreased by rapamycin, while stattic had no effect on mTOR activation. These results suggest that STAT3 functions downstream of mTORC1 signaling in tendon constructs.

The data suggest that the TGFβ1-mTORC1-STAT3 signaling axis plays a role in both physiological and pathological tendon conditions. TGFβ1 regulates tendon development and induces fibrotic scar formation in adult tendons. In tendon injury, TGFβ1 promotes fibroblast proliferation and reduces apoptosis during the proliferation phase of tendon healing [9]. mTORC1 signaling is critical role in tendon development and regeneration by regulating tenogenic differentiation [47]. Recent mouse genetic studies have reported that precise regulation of mTORC1 is necessary for proper tendon development and maturation [27]. STAT3 is activated during tendon healing to regulate inflammation in tendon stem and progenitor cells [48]. We demonstrated that mTORC1 and STAT3 partly mediate TGFβ1 function in tenogenic differentiation and matrix maturation. Together with other studies, current studies open new direction for investigating the TGFβ1-mTORC1-STAT3 signaling axis in tendon biology.

The current study has several limitations. First, we used small-molecule inhibitors, including rapamycin and stattic, for the rescue experiments. Although these are well-established inhibitors of mTORC1 and STAT3, they could non-specifically inhibit other signaling pathways. Future studies using genetic modifications specific to mTORC1 and STAT3 will help address this limitation. Fortunately, the 3D tendon constructs can be genetically modified using specific mouse tendon cells and the Adeno-Cre virus system, which will be a useful tool for future in vitro approaches. Second, the 3D tendon culture system is limited to fully capture the in vivo complexity of TGFβ1-mTORC1-STAT3 signaling in tendon tissue. Further genetic studies using mouse models will be a valuable next step. Finally, we observed only partial effects of the inhibitors on TGFβ1-induced 3D tendon formation, suggesting that additional signaling pathways are involved in TGFβ1 function in the 3D tendon formation. Elucidating other molecular and cellular mechanisms that mediate TGFβ1 function in tendon formation will be an exciting future direction. Overall, this in vitro pharmacological rescue experiment lays the groundwork for future molecular and genetic studies to further investigate the TGFβ1-mTORC1-STAT3 signaling axis in tendon biology.

## Acknowledgement

We would like to thank Penn Center for Musculoskeletal Disorders (PCMD) Histology and Biomechanics Core for technical assistance with the histology and 3D printing, respectively. We also thank Cell and Developmental Biology (CDB) Microscopy Core for technical assistance with confocal microscopy. This work was supported in part by a grant from the U.S. National Institutes of Health (NIH), National Institute of Arthritis and Musculoskeletal and Skin Diseases (KSJ: R01 AR079486). The PCMD Cores were supported by a grant from NIH/NIAMS P30AR069619.

## Author Contributions

Study conception and design: KSJ, BK; Acquisition of data: BK, AS; Analysis and interpretation of data: BK, AS; Drafting of the manuscript: BK, KSJ.

## Data Availability Statement

The data that support the findings of this study are available within the manuscript.

## References

1. Thorpe, C.T. and H.R. Screen, Tendon Structure and Composition. Adv Exp Med Biol, 2016. 920: p. 3–10.

2. Zhang, G., et al., Development of tendon structure and function: regulation of collagen fibrillogenesis. J Musculoskelet Neuronal Interact, 2005. 5(1): p. 5–21.

3. Andarawis-Puri, N., E.L. Flatow, and L.J. Soslowsky, Tendon basic science: Development, repair, regeneration, and healing. J Orthop Res, 2015. 33(6): p. 780–4.

4. Huang, A.H., H.H. Lu, and R. Schweitzer, Molecular regulation of tendon cell fate during development. J Orthop Res, 2015. 33(6): p. 800–12.

5. Murchison, N.D., et al., Regulation of tendon differentiation by scleraxis distinguishes force-transmitting tendons from muscle-anchoring tendons. Development, 2007. 134(14): p. 2697–708.

6. Schweitzer, R., et al., Analysis of the tendon cell fate using Scleraxis, a specific marker for tendons and ligaments. Development, 2001. 128(19): p. 3855–66.

7. Docheva, D., et al., Tenomodulin is necessary for tenocyte proliferation and tendon maturation. Molecular and Cellular Biology, 2005. 25(2): p. 699–705.

8. Li, Y., T. Wu, and S. Liu, Identification and Distinction of Tenocytes and Tendon-Derived Stem Cells. Front Cell Dev Biol, 2021. 9: p. 629515.

9. Li, Y., et al., Transforming growth factor-beta signalling pathway in tendon healing. Growth Factors, 2022. 40(3-4): p. 98–107.

10. Forslund, C., BMP treatment for improving tendon repair. Studies on rat and rabbit Achilles tendons. Acta Orthop Scand Suppl, 2003. 74(308): p. I, 1-30.

11. Li, X., et al., CTGF induces tenogenic differentiation and proliferation of adipose-derived stromal cells. J Orthop Res, 2019. 37(3): p. 574–582.

12. Disser, N.P., et al., Insulin-like growth factor 1 signaling in tenocytes is required for adult tendon growth. FASEB J, 2019. 33(11): p. 12680–12695.

13. Yao, L., et al., Phenotypic drift in human tenocyte culture. Tissue Eng, 2006. 12(7): p. 1843–9.

14. Nichols, A.E.C., et al., Impact of isolation method on cellular activation and presence of specific tendon cell subpopulations during in vitro culture. FASEB J, 2021. 35(7): p. e21733.

15. Wunderli, S.L., U. Blache, and J.G. Snedeker, Tendon explant models for physiologically relevant invitro study of tissue biology - a perspective. Connect Tissue Res, 2020. 61(3-4): p. 262–277.

16. Koo, B.H., et al., Characterization of TGFbeta1-induced tendon-like structure in the scaffold-free three-dimensional tendon cell culture system. Sci Rep, 2024. 14(1): p. 9495.

17. Yu, L., M.C. Hebert, and Y.E. Zhang, TGF-beta receptor-activated p38 MAP kinase mediates Smad-independent TGF-beta responses. EMBO J, 2002. 21(14): p. 3749–59.

18. Rodriguez-Garcia, A., et al., TGF-beta1 targets Smad, p38 MAPK, and PI3K/Akt signaling pathways to induce PFKFB3 gene expression and glycolysis in glioblastoma cells. FEBS J, 2017. 284(20): p. 3437-3454.

19. Kaji, D.A., et al., Tgfbeta signaling is required for tenocyte recruitment and functional neonatal tendon regeneration. Elife, 2020. 9.

20. Kallenbach, J.G., et al., Altered TGFB1 regulated pathways promote accelerated tendon healing in the superhealer MRL/MpJ mouse. Sci Rep, 2022. 12(1): p. 3026.

21. Gumucio, J.P., K.B. Sugg, and C.L. Mendias, TGF-beta superfamily signaling in muscle and tendon adaptation to resistance exercise. Exerc Sport Sci Rev, 2015. 43(2): p. 93–9.

22. Subramanian, A., et al., Mechanical force regulates tendon extracellular matrix organization and tenocyte morphogenesis through TGFbeta signaling. Elife, 2018. 7.

23. Heinemeier, K., et al., Role of TGF-beta1 in relation to exercise-induced type I collagen synthesis in human tendinous tissue. J Appl Physiol (1985), 2003. 95(6): p. 2390-7.

24. Saxton, R.A. and D.M. Sabatini, mTOR Signaling in Growth, Metabolism, and Disease. Cell, 2017. 168(6): p. 960–976.

25. Wang, Y. and J. Li, Current progress in growth factors and extracellular vesicles in tendon healing. Int Wound J, 2023. 20(9): p. 3871–3883.

26. Alenchery, R.G., et al., PAI-1 mediates TGF-beta1-induced myofibroblast activation in tenocytes via mTOR signaling. J Orthop Res, 2023. 41(10): p. 2163–2174.

27. Lim, J., et al., mTORC1 Signaling is a Critical Regulator of Postnatal Tendon Development. Scientific Reports, 2017. 7.

28. Nie, D.B., et al., Mechanical Overloading Induced-Activation of mTOR Signaling in Tendon Stem/Progenitor Cells Contributes to Tendinopathy Development. Frontiers in Cell and Developmental Biology, 2021. 9.

29. Hillmer, E.J., et al., STAT3 signaling in immunity. Cytokine Growth Factor Rev, 2016. 31: p. 1–15.

30. Bai, L.C., et al., A Potent and Selective Small-Molecule Degrader of STAT3 Achieves Complete Tumor Regression. Cancer Cell, 2019. 36(5): p. 498-+.

31. Nakama, K., et al., Interleukin-6-induced activation of signal transducer and activator of transcription-3 in ruptured rotator cuff tendon. Journal of International Medical Research, 2006. 34(6): p. 624–631.

32. Ray, S., et al., The I1-6 Trans-Signaling-STAT3 Pathway Mediates ECM and Cellular Proliferation in Fibroblasts from Hypertrophic Scar. Journal of Investigative Dermatology, 2013. 133(5): p. 1212–1220.

33. Rawlings, J.S., K.M. Rosler, and D.A. Harrison, The JAK/STAT signaling pathway. Journal of Cell Science, 2004. 117(8): p. 1281–1283.

34. Kim, J.H., M.S. Yoon, and J. Chen, Signal Transducer and Activator of Transcription 3 (STAT3) Mediates Amino Acid Inhibition of Insulin Signaling through Serine 727 Phosphorylation. Journal of Biological Chemistry, 2009. 284(51): p. 35425–35432.

35. Lui, P.P.Y. and C.M. Wong, Biology of Tendon Stem Cells and Tendon in Aging. Front Genet, 2019. 10: p. 1338.

36. Park, N.R., et al., Reticulocalbin 3 is involved in postnatal tendon development by regulating collagen fibrillogenesis and cellular maturation. Sci Rep, 2021. 11(1): p. 10868.

37. Gaut, L. and D. Duprez, Tendon development and diseases. Wiley Interdiscip Rev Dev Biol, 2016. 5(1): p. 5–23.

38. Maity, S., et al., TGFβ acts through PDGFRβ to activate mTORC1 via the Akt/PRAS40 axis and causes glomerular mesangial cell hypertrophy and matrix protein expression. Journal of Biological Chemistry, 2020. 295(42): p. 14262–14278.

39. Zhai, X.X., et al., Expression of TGF-β1/mTOR signaling pathway in pathological scar fibroblasts. Molecular Medicine Reports, 2017. 15(6): p. 3467–3472.

40. Cheng, K.Y. and M. Hao, Mammalian Target of Rapamycin (mTOR) Regulates Transforming Growth Factor-β(TGF-β)-Induced Epithelial-Mesenchymal Transition via Decreased Pyruvate Kinase M2 (PKM2) Expression in Cervical Cancer Cells. Medical Science Monitor, 2017. 23: p. 2017–2028.

41. Li, C., et al., Noncanonical STAT3 Activation Regulates Excess TGF-β1 and Collagen I Expression in Muscle of Stricturing Crohn’s Disease. Journal of Immunology, 2015. 194(7): p. 3422–3431.

42. Chakraborty, D., et al., Activation of STAT3 integrates common profibrotic pathways to promote fibroblast activation and tissue fibrosis (Oct, 10.1038/s41467-017-01236-6, 2017). Nature Communications, 2021. 12(1).

43. Tang, L.Y., et al., Transforming Growth Factor-β (TGF-β) Directly Activates the JAK1-STAT3 Axis to Induce Hepatic Fibrosis in Coordination with the SMAD Pathway. Journal of Biological Chemistry, 2017. 292(10): p. 4302–4312.

44. Vella, A., et al., mTOR and STAT3 Pathway Hyper-Activation is Associated with Elevated Interleukin-6 Levels in Patients with Shwachman-Diamond Syndrome: Further Evidence of Lymphoid Lineage Impairment. Cancers, 2020. 12(3).

45. Dodd, K.M., et al., mTORC1 drives HIF-1α and VEGF-A signalling via multiple mechanisms involving 4E-BP1, S6K1 and STAT3. Oncogene, 2015. 34(17): p. 2239-2250.

46. Yokogami, K., et al., Serine phosphorylation and maximal activation of STAT3 during CNTF signaling is mediated by the rapamycin target mTOR. Current Biology, 2000. 10(1): p. 47–50.

47. Cong, X.X., et al., Activation of AKT-mTOR Signaling Directs Tenogenesis of Mesenchymal Stem Cells. Stem Cells, 2018. 36(4): p. 527–539.

48. Tarafder, S., et al., Tendon stem/progenitor cells regulate inflammation in tendon healing via JNK and STAT3 signaling. Faseb Journal, 2017. 31(9): p. 3991–3998.

